# Commensal bacteria promote type I interferon signaling to maintain immune tolerance

**DOI:** 10.1101/2021.10.21.464743

**Authors:** Adriana Vasquez Ayala, Chia-Yun Hsu, Kazuhiko Matsuo, Ekaterina Buzun, Marvic Carrillo Terrazas, Luke R. Loomis, Hsueh-Han Lu, Jong Hwee Park, Paul Rivaud, Matt Thomson, Hiutung Chu

## Abstract

Type I interferons (IFN) exert a broad range of biological effects important in coordinating immune responses. Host and microbial factors regulate IFN production, triggering a signaling cascade that has classically been studied in the context of pathogen clearance. In particular, commensal bacteria have been shown to induce IFN to protect against viral infections. Yet, whether immunomodulatory bacteria operate through IFN pathways to support immune tolerance remains elusive. Here, we demonstrate microbiota-dependent IFN signaling is required for priming tolerogenic T regulatory cells (Tregs) by intestinal dendritic cells (DCs). DCs deficient in IFN signaling through deletion of IFNAR-1 display dysregulated cytokine production in response to the commensal bacteria *Bacteroides fragilis*, resulting in blunted downstream Treg responses. Single cell RNA sequencing of gut tissues demonstrated that colonization with *B. fragilis* promotes a distinct type I IFN gene signature in Tregs during homeostasis and intestinal inflammation. Moreover, *B. fragilis-*mediated protection during experimental colitis was abrogated in IFNAR1-deficient mice. Altogether, our findings demonstrate an important role of microbiota-mediated immune tolerance via tonic type I IFN signaling.

## INTRODUCTION

Type I interferons (IFN) are involved in many essential immune functions, influencing both innate and adaptive immune effector responses (Trinchieri, 2010; Ivashkiv and Donlin, 2014; McNab et al., 2015). Type I IFNs, namely IFNα and IFNβ, are produced upon sensing microbial products resulting in the expression of interferon-stimulated genes (ISGs). While several hundred ISGs with various known functions have been identified, type I IFN has been primarily studied in its role in antiviral immunity (Schoggins et al., 2011; Schoggins, 2019). Recent work revealed that commensal microbes are involved in maintaining tonic type I IFN necessary to mount an effective antiviral immune response (Abt et al., 2012; Ganal et al., 2012; Yang et al.; Bradley et al., 2019; Wirusanti et al., 2022). Apart from the induction of antiviral ISGs, type I IFN can promote dendritic cell (DC) activation and maturation to enhance antigen presentation to prime adaptive (Honda et al., 2003; Simmons et al., 2012). Further, tonic type I IFN expression is essential for effective T cell responses (Aman et al., 1996; Levings et al., 2001; Bilsborough et al., 2003; Teijaro et al., 2013; Wilson et al., 2013). These studies highlight a potential role for microbial-induced IFN signaling in host immunity beyond antiviral responses. Of particular interest is the divergent effect of type I IFN on immune responses that depend on the context of microbial exposure. Type I IFN responses to microbial pathogens generate a robust antimicrobial and pro-inflammatory response via activation of ISGs (McNab et al., 2015; Boxx and Cheng, 2016). In contrast, detection of commensal products during homeostatic conditions triggers type I IFN signaling to support anti-inflammatory responses (Lee and Ashkar, 2018). While this tonic IFN expression is critical in priming immune cells to rapidly mobilize (Abt et al., 2012; Ganal et al., 2012; Yang et al.), it also promotes tolerance by suppressing proliferation of pathogenic T cell responses and the production of pro-inflammatory cytokines. Previous studies also suggest that type I IFN may influence regulatory T cell (Treg) function. Notably, signaling via interferon-α/β receptor 1 (IFNAR1) is required for Treg expansion and suppression of pathogenic T cells during colitis (Lee et al., 2012; Stewart et al., 2013; Kawano et al., 2018). In humans, several genes in the IFN pathway have been associated with inflammatory bowel disease (IBD) susceptibility in genome wide association studies. *IFNAR1* is suggested to be an IBD risk allele, and single nucleotide polymorphisms (SNPs) in *JAK2, TYK2, STAT1*, and *STAT3* have major consequences in the JAK/STAT pathway, including dysregulation of cytokine production (Jostins et al., 2012). These findings implicate the potential for type I IFN in the maintenance of mucosal homeostasis and immune tolerance.

Clinical and experimental evidence has implicated the gut microbiota in governing host immunity during steady-state and disease (Hooper et al., 2012; Belkaid and Hand, 2014). In particular, the mechanism by which commensal bacteria regulate type I IFN responses to maintain mucosal immunity is of significant interest. Microbiota-induced IFN pathways have been shown to be critical in mounting antiviral resistance in the lung (Steed et al., 2017; Bradley et al., 2019). Moreover, glycolipids from commensal *Bacteroides* have been reported to direct antiviral function via IFNβ expression in DCs (Stefan et al., 2020). These studies indicate the microbiota regulate type I IFN responses to influence adaptive immunity including DC and T cell function. Furthermore, published studies have demonstrated type I IFN is involved in the maintenance of Tregs in the gut (Lee et al., 2012; Kole et al., 2013; Nakahashi-Oda et al., 2016). Given the role of the microbiota in maintaining tonic IFN expression and our work with Treg-promoting commensal bacteria, we hypothesize that immunomodulatory bacteria signal via type I IFN pathways to direct intestinal immune tolerance.

Here, we reveal that tonic type I IFN is maintained by commensal bacteria and required for tolerogenic immune responses in the gut. Previous work from our group and others demonstrated that *Bacteroides fragilis* prime DCs to promote Foxp3+ Treg responses to control intestinal inflammation (Shen et al., 2012; Chu et al., 2016). In the present study, we expand these finding by demonstrating that germ-free (GF) mice are indeed deficient in IFN responses, and colonization with a single commensal bacterium, *B. fragilis*, partially restores tonic type I IFN in the gut comparable to specific pathogen free (SPF) mice. We also reveal select commensal bacteria demonstrate variable induction of IFN signaling in DCs, suggesting this immunomodulatory trait is unique among certain microbes. Notably, *B. fragilis*-induced IFN in DCs are required for commensal-induced Foxp3+ Treg responses. Indeed, while investigating *B. fragilis*-mediated gene signatures in Treg cells, we discovered an enrichment of IFN related genes among intestinal Foxp3+ Treg cells. Finally, this commensal-driven IFN signaling in gut DCs is necessary for protection from experimental colitis. Our findings demonstrate that commensal bacteria promote intestinal homeostasis through type I IFN signaling.

## RESULTS

### Commensal bacteria direct intestinal type I IFN responses

Emerging evidence suggests commensal bacteria are important regulators of tonic type I IFN signaling (Sonnenburg et al., 2006; Yamamoto et al., 2012; Schaupp et al., 2020; Di Domizio et al., 2020; Lam et al., 2021) and are required to mount an effective immune response to pathogens (Abt et al., 2012; Steed et al., 2017; Bradley et al., 2019; Winkler et al., 2020; Erttmann et al., 2022). Given the pleiotropic effects of type I IFN, we examined the association between the microbiota and type I IFN signaling in the gut during steady state. Expression of IFN-related genes was assessed in colon tissue of GF and SPF C57BL/6J mice. The presence of a commensal microbial community was required for the induction of *Ifnb* and *Mx1* (an IFN stimulated gene) expression in the colon (Figure 1A). Particularly, mono-colonization of GF mice with the commensal bacterium, *B. fragilis*, was able to partially restore IFN-related gene expression in the colon (Figure 1A). We next investigated the contribution of the microbiota in priming local intestinal type I IFN responses upon microbial stimulation. Colon explants from GF and SPF mice were treated with polyinosinic-polycytidylic acid (poly I:C), a Toll-like receptor 3 (TLR-3) agonist and potent inducer of IFNβ. While poly I:C induced a significant increase in IFNβ production in SPF colon explants, GF tissues remained unresponsive to poly I:C (Figure 1B). Since type I IFN are constitutively expressed in the intestines by CD11c+ DCs (Kole et al., 2013; Schaupp et al., 2020; Stefan et al., 2020), we next investigated whether the presence of commensal bacteria influenced IFNα/β receptor (IFNAR)-dependent signaling in gut DCs. Colonic lamina propria (cLP) cells were isolated from SPF and GF mice and stimulated with recombinant IFNβ ex vivo. Colonic CD11c+ DCs from SPF mice responded to IFNβ stimulation via increased phosphorylation of signal transducer and activator of transcription 1 (STAT1), as measured by flow cytometry (Figure 1C). However, pSTAT1 expression remained unchanged in colonic CD11c+ cells from GF mice following IFNβ stimulation, suggesting impaired IFNAR signaling in mice lacking commensal bacteria. To investigate whether this IFN defect in the intestinal environment extends to systemic compartments, splenocytes from GF and SPF mice were treated ex vivo with poly I:C. As expected, at baseline SPF splenocytes expressed higher levels of type I IFN, as well as other cytokines and chemokines, compared to GF (Figure S1). Moreover, SPF splenocytes were more responsive to poly I:C stimulation than treated GF splenocytes. No induction of IFNα and IFNɣ (a type II IFN) were observed upon poly I:C treatment of GF splenocytes. Further, while poly I:C induced IFNβ in GF splenocytes, expression was limited and equivalent to untreated SPF cells (Figure S1). To confirm the requirement of commensal bacteria for type I IFN response in the intestinal environment in vivo, GF, *B. fragilis* mono-colonized, and SPF mice were treated with poly I:C by intraperitoneal (IP) injection and evaluated for IFN responsiveness. SPF mice treated with poly I:C demonstrated significant expression of type I IFN related genes (Figure 1D-1F and Figure S2). In contrast, GF mice injected with poly I:C showed no response in comparison to the PBS control 4 hours post-injection (Figure 1D-G and Figure S2), consistent with ex vivo studies (Figure 1A-C). To verify whether this IFN defect in GF mice can be restored with commensal bacteria, we colonized GF mice with *B. fragilis*. Indeed, poly I:C treatment of *B. fragilis* mice led to a significant induction of IFNβ and IFN related genes, indicating the presence of commensal bacteria is sufficient to restore homeostatic type I IFN responses (Figure 1D-G and Figure S2). These data establish that commensal bacteria are critical in maintaining type I IFN responses in intestinal tissues.

**Figure 1.**
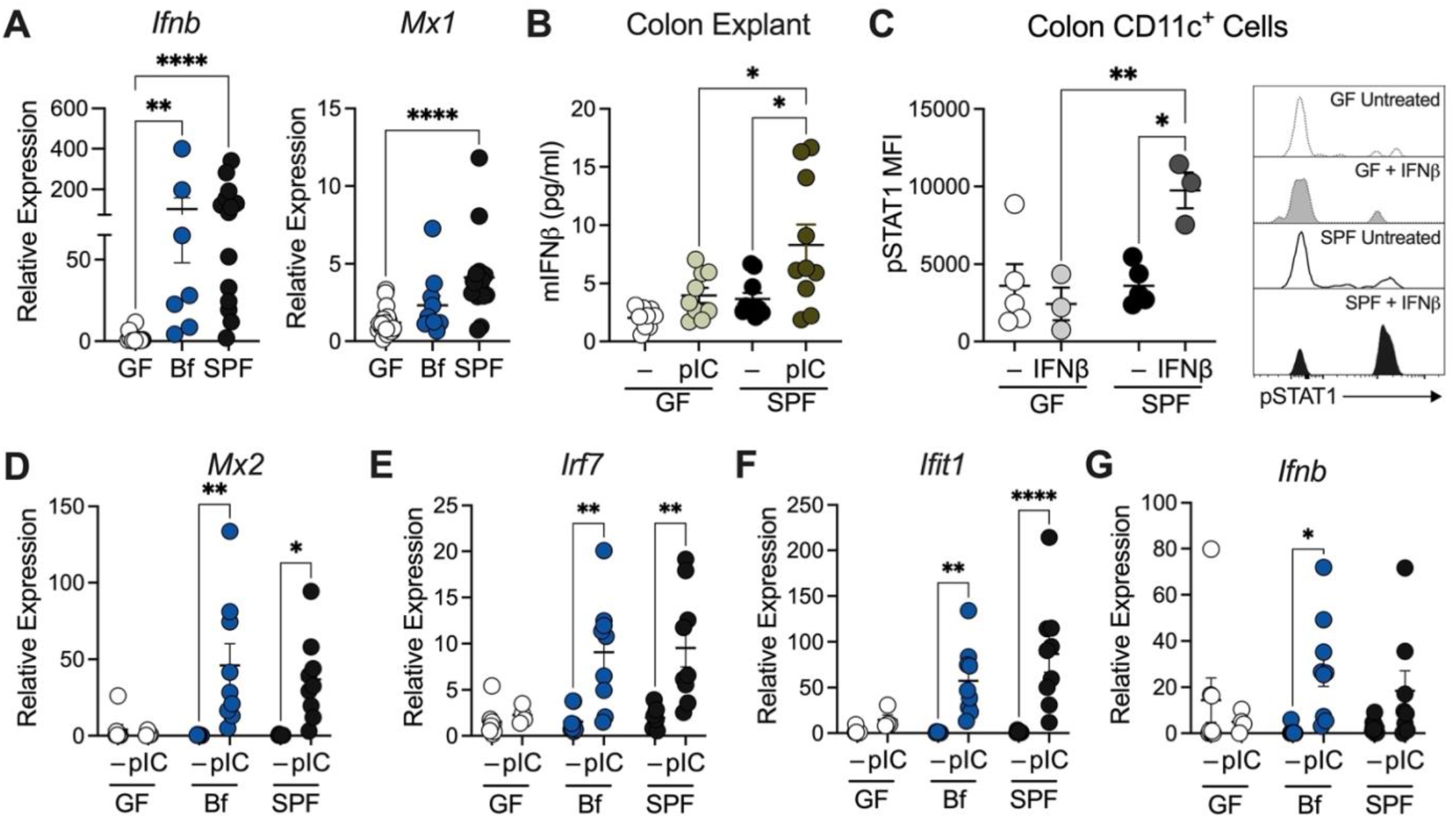
Commensal bacteria maintain intestinal type I IFN responses. (**A**) Expression of *Ifnb* and *Mx1* in colon tissue from germ-free (GF), *B. fragilis* mono-colonized (Bf), and specific pathogen-free (SPF) as measured by RT-qPCR. Each point represents a single mouse. (**B**) GF and SPF colon explants were cultured with or without stimulation of poly I:C (pIC; 2 μg/ml) for 24 hours. Supernatant was then collected and measured for IFNβ secretion by ELISA. Each point represents a single mouse. (**C**) Colonic lamina propria cells were isolated from GF and SPF mice and stimulated with IFNβ (25 ng/ml). pSTAT1 was assessed by flow cytometry. Each point represents colons pooled from multiple mice, with n=10 per group. (**D-G**) GF, *B. fragilis* mono-colonized (Bf), and SPF mice were injected (IP) with 100 µg/ml poly I:C (pIC) and colon tissues were harvested after 4 h post-injection. Gene expression for **(D)** *Mx2*, **(E)** *Irf7*, **(F)** *Ifit1*, **(G)** *Ifnb* was measured. Each point represents a single mouse. Data are representative of at least two independent experiments. Statistical significance was determined by Kruskal-Wallis, unpaired t-test, and 2way ANOVA. p<0.05 (*), p<0.01 (**), p<0.001 (***), and p<0.0001 (****).

### Select commensal bacteria induce IFN signaling in dendritic cells

Dendritic cells establish and maintain the local gut immune environment by sampling luminal contents, including bacteria (Rescigno and Sabatino, 2009). Furthermore, we and others have demonstrated that DCs orchestrate *B. fragilis*-mediated immune tolerance via bacterial sensing through pattern recognition receptors (Shen et al., 2012; Dasgupta et al., 2014; Chu et al., 2016). Additionally, commensal bacteria concurrently trigger these pattern recognition receptors to produce type I IFN, revealing gut DCs as a source of type I IFN (Fitzgerald-Bocarsly and Feng, 2007; Kawashima et al., 2013). This evidence led us to hypothesize that *B. fragilis* facilitates tolerogenic responses in the gut by engaging IFNAR signaling in DCs. To evaluate if *B. fragilis-*mediated immune tolerance is, in part, achieved via IFN, we measured IFN related gene expression in DCs treated with *B. fragilis* in vitro. Treatment of bone marrow derived dendritic cells (BMDCs) with *B. fragilis* induced expression and secretion of IFNβ, but not IFNα or IFNɣ (Figures 2A and 2B), consistent with in vivo findings (Figures 1D-1G). Next, we leveraged the *IFNb*^mob^ EYFP reporter mouse to examine *B. fragilis* induced IFNβ among conventional and plasmacytoid DCs (Figure 2C). Treatment with *B. fragilis* induced IFNβ-YFP expression in both DCs subsets. We then examined whether colonization with *B. fragilis* induced IFN expression among colonic DCs. To capture the levels of IFNβ contributed by intestinal DCs primed by *B. fragilis* in vivo, colonic CD11c+ cells were isolated from GF and *B. fragilis* mono-colonized mice and IFNβ secretion was measured at baseline (without exogenous treatment). Indeed, colonic lamina propria CD11c+ cells from *B. fragilis* mono-colonized mice produced higher levels of IFNβ compared to GF mice (Figure 2D). To test if *B. fragilis* stimulates IFN signaling in DCs, pSTAT1 levels were measured by flow cytometry. Expression of pSTAT1 in BMDCs was significantly increased in response to *B. fragilis* treatment (Figure 2E), consistent with induction of autocrine IFN signaling. As expected, this increase was lost in *Ifnar1*^-/-^ DCs (Figure S3A). We next examined whether other commensal bacteria were similarly able to induce type I IFN responses as observed for *B. fragilis*. Several studies have also reported on specific bacterial strains capable of inducing IFNβ production, including *Bacteroides* species (Steed et al., 2017; Stefan et al., 2020; Winkler et al., 2020; Yang et al.). Notably, *Escherichia coli* (Ec) and *Bifidobacterium longum* (Bl) were able to induce pSTAT1 among DCs at levels comparable to *B. fragilis* (Figure 2F). In contrast, other commensal bacteria tested did not support significant pSTAT1 induction among DCs (Figure 2F). Due to the selectivity in pSTAT1 induction between *Bacteroides* species we next measured IFNβ production in the supernatant of commensal treated-DCs. Treatment with *Bacteroides thetaiotaomicron* (Bt) and *Bacteroides vulgatus* (Bv) led to significant IFNβ secretion, at comparable levels to *B. fragilis* (Figure 2G). Our findings here suggest variation in bacterial-driven priming of IFN responses amongst commensals in the gut, including distinct properties in signaling via pSTAT1 (Figure 2F) versus IFNβ production (Figure 2G and S3B). Of note, STAT1 and IFN signaling have been implicated in promoting both gut homeostasis and driving intestinal inflammation (Cho and Kelsall, 2014; Giles et al., 2017). One potential explanation for this divergent effect may be that both type I (IFNα/β) and type II (IFNɣ) IFN share many of the same signaling components, including STAT1. As such, commensal and pathogenic microbes, signaling through distinct subset of pattern recognition receptors, will calibrate different levels of type I and type II IFNs that ultimately will drive either a tolerogenic or inflammatory immune response. Altogether, select commensals, including *B. fragilis*, drive tonic type I IFN production and promote autocrine IFN signaling in DCs.

**Figure 2.**
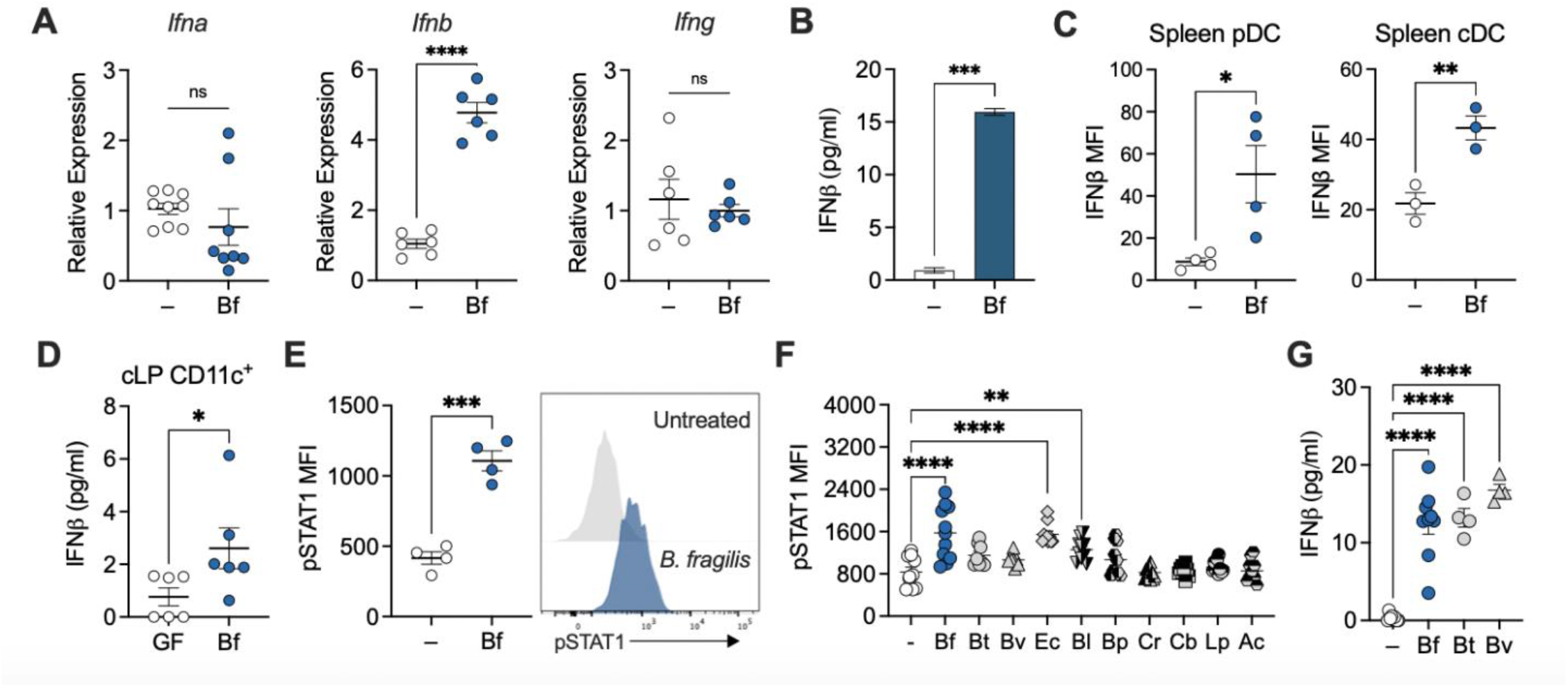
*B. fragilis* induces IFNβ expression in dendritic cells to coordinate Treg response. Bone marrow derived dendritic cells (BMDCs) were treated with *B. fragilis* for 18 hours and **(A)** relative expression of *Ifna, Ifnb*, and *Ifng* were measured by qRT-PCR. (**B**) Supernatants from BMDC cultures were collected and IFNβ secretion was measured by ELISA. (**C**) Splenocytes were isolated from IFNβ reporter mice and MFI of IFNβ was quantified by flow cytometry in cDCs and pDCs. (**D**) Colon lamina propria cells were isolated and enriched for CD11c+ cells from GF and Bf mono-colonized mice and cultured for 18 hours. Supernatant was collected and measured for IFNβ by ELISA. Each point represents colons pooled from five mice per point and represent n=30 for each group. (**E**) BMDCs from SPF mice were treated with *B. fragilis* for 18 hours, and cells were stained for pSTAT1 and analyzed by flow cytometry. (**F**) BMDCs from SPF mice were treated with *B. fragilis, B. thetaiotaomicron, B. vulgatus, Escherichia coli, Bifidobacterium longum, Blautia producta, Clostridium ramosum, Clostridium butyricum, Lactobacillus plantarum*, and *Anaerostipes caccae* for 18 hours. Cells were stained and pSTAT1 was measured by flow cytometry. (**G**) BMDC supernatant from *B. fragilis, B. thetaiotaomicron*, and *B. vulgatus* were collected and measured for IFNβ by ELISA. Data are representative of at least two independent experiments. Statistical analysis was determined by unpaired t-test and two-way ANOVA. p<0.05 (*), p<0.01 (**), p<0.001 (***), and p<0.0001 (****).

### *B. fragilis* induces type I IFN expression in dendritic cells to coordinate Treg responses

It has been well documented that *B. fragilis* primes DCs to foster immune tolerance by inducing IL-10 expression in Tregs (Shen et al., 2012; Chu et al., 2016). Thus, we evaluated the necessity of type I IFN signaling for *B. fragilis*-induced immune tolerance using IFNAR1-deficient DCs. While *B. fragilis* significantly induced type I IFN related genes in wildtype (WT) DCs (e.g., *Ifnb, Mx1, Oas1*, and *Mx2*), *Ifnar1*^-/-^ DCs remained unresponsive to *B. fragilis* (Figure 3A and S4A). This induction appeared to be selective, as the expression of certain IFN-responsive genes (e.g., *Irf3, Irf9*) was either unaffected or inhibited by *B. fragilis* (Figure S4B and S4C). Additionally, expression of *Ifnα* was also unaffected by *B. fragilis* in WT DCs (Figure 2A), suggesting perhaps some specificity in the induction of type I IFN gene expression downstream of pattern recognition receptor signaling. Previous studies with *B. fragilis* demonstrated that defects in TLR2 (Round et al., 2011; Shen et al., 2012) and NOD2 (Chu et al., 2016) signaling led to impaired Treg responses. Here, we observed *B. fragilis*-induced pSTAT1 responses and IFNβ production required signaling via NOD2, a cytosolic pattern recognition receptor that recognizes a component of peptidoglycan (Figures S5A and S5B). In contrast, TLR2 signaling appears to be dispensable for *B. fragilis*-induced IFN responses, with increased pSTAT1 and IFNβ levels in *B. fragilis*-treated *Tlr2*^-/-^ DCs relative to WT DCs (Figures S5C and S5D). We also previously reported a shift in the cytokine milieu in DCs defective in microbial sensing, demonstrating a significant decrease in expression of IL-10, which is required for induction of tolerogenic Tregs (Chu et al., 2016). Similarly, here we observed a significant reduction in the anti-inflammatory cytokines IL-10 and IL-27 in *Ifnar1*^-/-^ BMDCs compared to *B. fragilis*-treated WT BMDCs (Figure 3B-3D). IL-27 is an immunoregulatory cytokine that targets Foxp3+ Tregs to mediate anti-inflammatory activity (Kim et al., 2019; Nguyen et al., 2019) while suppressing IL17-producing CD4+ T cells (Batten et al., 2006). This reduction in anti-inflammatory response is paired with a shift towards a pro-inflammatory environment, with an increase in IL-1β upon *B. fragilis* treatment of *Ifnar1*^-/-^ BMDCs (Figure 3E and 3F). *B. fragilis* treatment of BMDCs did not alter expression of pro-inflammatory IFNɣ, indicating selectivity in cytokine regulation by commensal microbes (Figure 3G). Consistent with these observations, we report extensive dysregulation of cytokine and chemokine production in *B. fragilis*-treated *Ifnar1*^-/-^ BMDCs (Figure S4D-S4J). These data support a critical role for IFN signaling in DCs to constrain pro-inflammatory responses upon sensing commensal bacteria.

**Figure 3.**
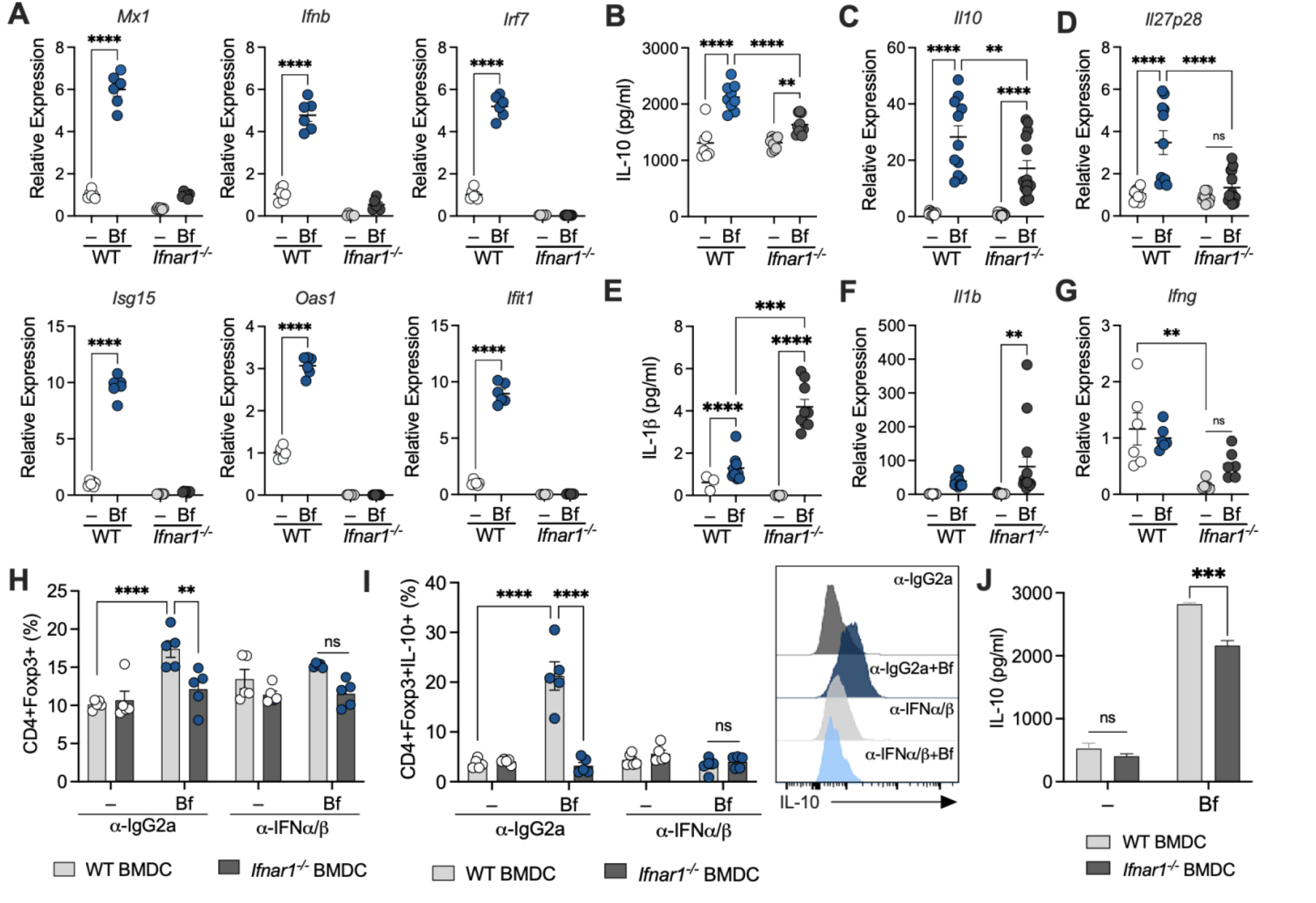
Type I IFN signaling in DCs are required to promote Treg responses. **(A-G)** WT and IFNAR-deficient BMDCs were pulsed with *B. fragilis* (Bf) for 18 hours. Cells were harvested and analyzed by qRT-PCR for gene expression of **(A)** IFN related genes, **(C)** *Il10*, **(D)** *Il27p28*, **(F)** *Il1b*, and **(G)** *Ifng*. **(B**,**E)** Supernatant from BMDC cultures were collected and cytokine secretion was measured for **(B)** IL-10 and **(E)** IL-1β by ELISA. **(H-I)** WT and IFNAR1-deficient BMDCs were untreated (–) or treated with *B. fragilis* (Bf) co-cultured with WT T-cells. Cells were treated with 10 µg/ml of IFNα/β neutralizing antibodies or isotype control (IgG2a) every 24 hours and then stained and analyzed by flow cytometry for **(H)** Foxp3+ Tregs and **(I)** IL-10 producing Foxp3+ Tregs and **(J)** supernatant were collected to measure IL-10 secretion by ELISA. Data are representative of at least two independent experiments. Statistical significance was determined by two-way ANOVA. p<0.05 (*), p<0.01 (**), p<0.001 (***), and p<0.0001 (****).

We hypothesized that this skewed pro-inflammatory environment driven by IFNAR-deficient DCs would restrain downstream Treg function. To test this hypothesis, WT and *Ifnar1*^-/-^ BMDCs were treated with *B. fragilis* and co-cultured with WT CD4+ T cells. Indeed, *B. fragilis* pulsed WT DCs supported the induction of CD4+ Foxp3+ Tregs (Figure 3H) and production of IL-10 (Figure 3I and 3J). However, *B. fragilis* treated *Ifnar1*^-/-^ BMDCs resulted in a significant reduction of IL-10+ Treg populations, indicating IFN signaling in DCs can impact subsequent T cell function despite intact IFNAR signaling in these WT CD4+ T cells. *B. fragilis*-treated *Ifnar1*^-/-^ DCs co-cultured with T cells also displayed decreased IL-10 secretion, as detected by ELISA, relative to WT DC:T cell cultures (Figure 3J). We next assessed whether neutralization of type I IFN phenocopies the Treg defect observed in *B. fragilis*-treated *Ifnar1*^-/-^ DCs. The induction of Treg and IL10 production by *B. fragilis* was abrogated upon addition of neutralizing antibodies for IFNα and IFNβ in DC:T cell co-cultures compared to isotype controls (Figure 3H and 3I). This suggests IFNα and/or IFNβ function as critical signals that, in their absence, impede development of *B. fragilis*-induced Tregs. Altogether, these data reveal type I IFN signaling in DCs is critical for commensal bacteria to direct appropriate Treg responses.

### IFN signaling is required for *B. fragilis*-mediated protection during colitis

Given the requirement of type I IFN signaling for *B. fragilis* induced Treg responses in vitro, we next asked whether the gene signature of intestinal Tregs is associated with tonic IFN upon bacterial colonization. To examine this link, we performed single cell RNA sequencing of gut Tregs from GF and *B. fragilis* mono-colonized mice. Tregs from mesenteric lymph nodes (MLN) of GF and *B. fragilis* mono-colonized mice partitioned into two distinct clusters (Figure 4A). Notably, Tregs from MLN and the colonic lamina propria (cLP) of *B. fragilis* mono-colonized mice exhibited a significant upregulation in IFN related genes, including *Ifit3, Isg15, Mx1, Stat1*, and *Oas* gene family (Figure 4B). It has been reported that suppressor of cytokine signaling 1 (SOCS1 and USP18, ubiquitin-specific peptidase 18), both potent negative regulators of type I IFN signaling, play a role in maintaining Treg differentiation and function (Takahashi et al., 2011, 2017; Yang et al., 2020). To investigate the relevance of this negative regulator in the context of the intestinal environment, we assessed the expression of *Socs1* and found expression was significantly upregulated in tissue from *B. fragilis* mono-colonized and SPF mice compared to GF (Figure S6A), which corroborates the induction of *Socs1* we observed in colonic Tregs of *B. fragilis* mono-colonized mice (Figure 4B). Expression of *Usp18* in whole colon tissue showed no significant differences among GF, *B. fragilis* mono-colonized, and SPF mice (Figure S6B), in contrast to the marked difference seen in Tregs from MLN and colon between GF and *B. fragilis* mono-colonized mice (Figure 4B). This suggests USP18 may possess distinct expression in Tregs in the intestine. These data support the notion that type I IFN activity is a prominent pathway induced in Tregs by commensal bacteria.

**Figure 4.**
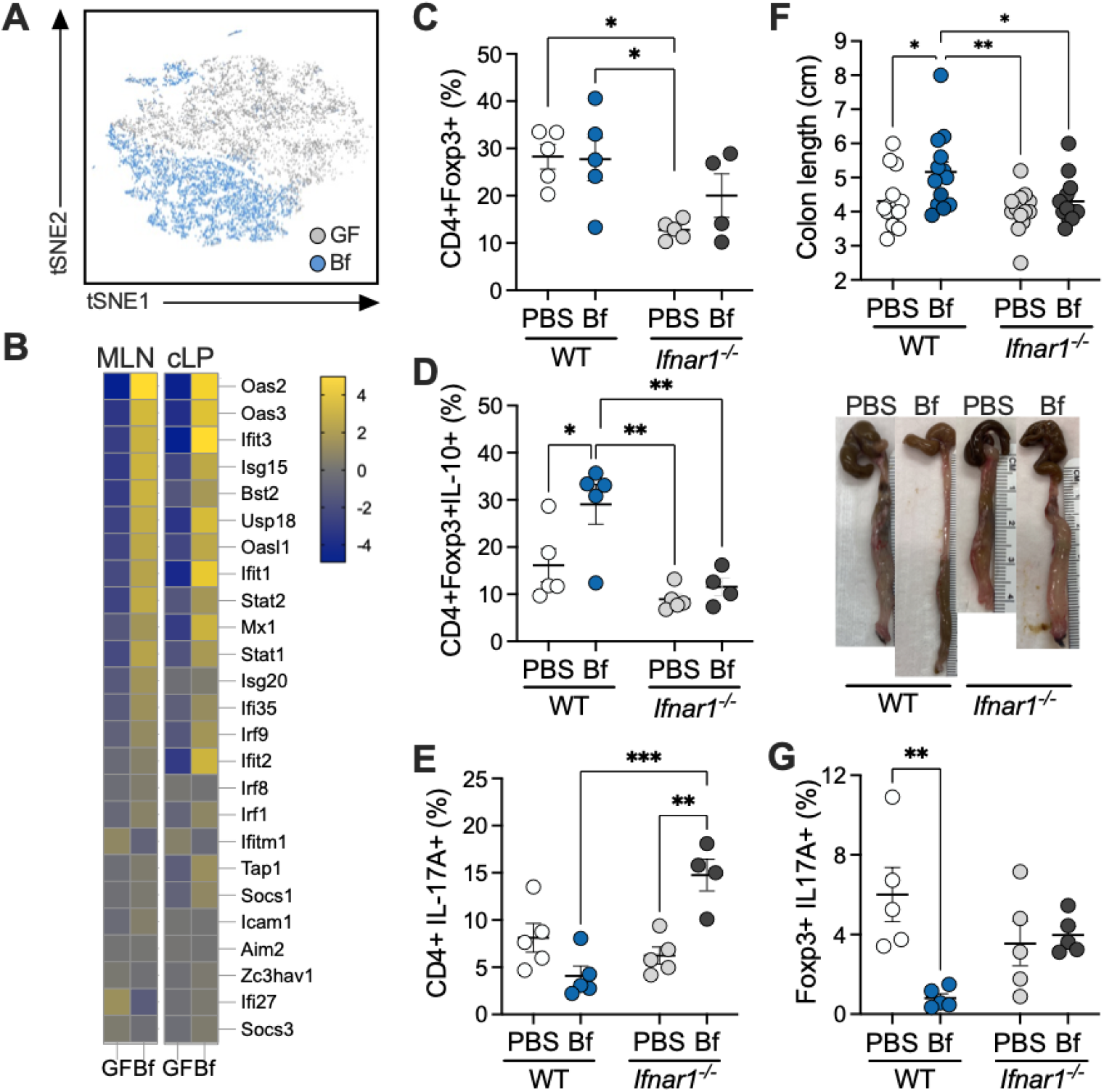
Bacterial colonization induces type I IFN signature in intestinal Tregs. **(A)** 4- to 6-week-old female germ-free (GF) Foxp3^hCD2^/IL-10^Venus^ mice were orally gavaged with a single dose of *B. fragilis* (10^8^ CFU resuspended in sterile PBS). Four weeks later, mesenteric lymph nodes (MLN) and colon tissues were harvested, and single suspensions were prepared. CD4+hCD2+ cells were enriched and subjected to single-cell RNA sequence analysis. t-SNE plot of Tregs from MLN using K-nearest neighbor graph-based clustering. Individual cells were colored by experimental groups. **(B)** Heatmap showing type I IFN signature genes from Tregs in MLN and colonic lamina propria (cLP). Genes are colored by the difference in log_2_-fold change expression between GF and *B. fragilis* mono-colonized mice. **(C-E) WT** and IFNAR1-deficient mice were gavaged with either PBS or *B. fragilis* for two week and proportions of **(C)** CD4+ Foxp3+ Tregs, **(D)** CD4+ Foxp3+ IL10+ Tregs, and **(E)** CD4+ Foxp3-IL17A+ T cells in cLP were quantified from lymphocytes isolated from the cLP. **(F-G)** WT and IFNAR1-deficient mice were orally treated with sterile PBS or *B. fragilis* during DNBS colitis.**(F)** Colon length after DNBS colitis and gross morphology of colon tissues. **(G)** Proportions of CD4+ Foxp3+ IL17A+ T cells in cLP after DNBS colitis. Each point represents a single mouse. Statistical significance was determined by two-way ANOVA. p<0.05 (*), p<0.01 (**), p<0.001 (***), and p<0.0001 (****).

Due to the striking interferon gene signature in *B. fragilis* mono-colonized mice, we investigated the ability of *B. fragilis* to direct Treg responses in WT and IFNAR-deficient SPF mice. As expected, oral administration of *B. fragilis* induced IL-10-producing Foxp3+ Tregs in WT SPF mice compared to PBS-treated controls (Figure 4D). However, *B. fragilis* induction of Treg responses was abolished in *Ifnar1*^-/-^ SPF mice (Figure 4D). Moreover, proportions of Foxp3+ Tregs were significantly reduced amongst *Ifnar1*^-/-^ mice (Figure 4C), consistent with previous work that reports loss of Treg induction in *Rag1*^*-/-*^ recipients upon transfer of *Ifnar1*^*-/-*^ Tregs (Lee et al., 2012). Dysregulation of Treg functions is associated with increased frequency of pathogenic T cell subsets such as Th1 or Th17. Since Treg populations are significantly reduced in *Ifnar1*^-/-^ mice, we investigated whether Th17 subsets were reciprocally elevated due to this imbalance. Indeed, we observed a significant induction of CD4+IL17A+ T cells in *Ifnar1*^-/-^ mice treated with *B. fragilis*, compared to WT counterparts (Figure 4E), indicating a critical role of type I IFN signaling in maintaining Treg/Th17 balance. We next examined whether type I IFN responses are essential for immune tolerance during colitis. WT or *Ifnar1*^-/-^ mice were orally treated with *B. fragilis* during chemically-induced colitis (DNBS, dinitrobenzenesulfonic acid). *B. fragilis* administration ameliorates DNBS colitis in WT mice, as previously reported (Shen et al., 2012; Chu et al., 2016). However, protection by *B. fragilis* was lost in *Ifnar1*^-/-^ groups, as indicated by shortened colons and thickening of the mucosa (Figure 4F). We previously demonstrated that *B. fragilis*-induced Foxp3+ Tregs were required for protection against DNBS colitis. We thus examined populations of Foxp3+ Tregs and its production of IL-10, along with Th1 and Th17 subsets. While we observed some trends, no significant differences in these cell subsets were detected in WT and *Ifnar1*^-/-^ mice during experimental colitis (Figure S7A-S7H). However, cLP Foxp3+ Tregs isolated from *Ifnar1*^-/-^ mice following induction of DNBS colitis revealed increased IL-17A production, relative to WT controls treated with *B. fragilis*. In parallel to increased IL-17A production in the colon, elevated levels of RORgt+ IL-17A+ populations were observed in the MLNs of *Ifnar1*^-/-^ mice during DNBS colitis (Figure S7H). These findings indicate that in the absence of commensal-mediated IFN signaling, Tregs adopt a more inflammatory signature. Altogether, this study establishes the requirement of commensal-driven type I IFN signaling in Treg function and highlights the pro-inflammatory shift that occurs in a type I IFN-deficient intestinal environment.

## DISCUSSION

Efforts to understand how the microbiota primes the IFN system have largely focused on its involvement in anti-viral immunity (Steed et al., 2017; Bradley et al., 2019; Schaupp et al., 2020; Stefan et al., 2020; Yang et al.; Erttmann et al., 2022). The role of commensal-induced IFN in promoting intestinal tolerance, however, remains elusive. Several studies have suggested type I IFN and STAT1 (the main transducer of IFN) may be involved in mediating immune tolerance. Type I IFN and STAT1 signaling have been reported to drive expansion of Tregs and promote expression of IL-10, while suppressing intestinal T cells producing IL-17 (Guo et al., 2008; Lee et al., 2012; Stewart et al., 2013; Kole et al., 2013). In contrast, other reports indicate STAT1 and type I IFN induced during viral infections inhibits Treg cell proliferation and activation of Foxp3+ Tregs (Ma et al., 2011; Srivastava et al., 2014a, 2014b). These discordant findings highlight the duality of type I IFN responses, capable of driving both immunostimulatory and suppressive regulatory functions. Further, these studies indicate the context in which microbes induce IFN may determine whether proinflammatory or tolerogenic immune responses are elicited. We now present findings that support a model whereby commensal bacteria drive autocrine IFN signaling in DCs during steady state conditions, which is essential for directing Treg development and promoting mucosal homeostasis.

Mounting evidence indicates that intestinal DCs produce and maintain type I IFN in response to the gut microbiota, and microbiota depletion results in severely diminished type I IFN production (Kole et al., 2013; Kawashima et al., 2013; Stefan et al., 2020). Recently, several studies demonstrated that commensal bacterium *B. fragilis* induces IFNβ among antigen presenting cells to trigger systemic anti-viral responses (Erttmann et al., 2022; Stefan et al., 2020). Accordingly, we discovered that select human commensal bacteria, including *B. fragilis*, are capable of driving IFNβ production and pSTAT1 signaling in colonic and bone marrow derived DCs. We further expand these findings by uncovering an underappreciated role of commensal-induced IFN signaling to establish immune tolerance. Mechanistically, we demonstrate that autocrine IFNAR signaling in DCs is important in directing anti-inflammatory responses, as *Ifnar1*^-/-^ DCs and IFNα/β neutralization studies reveal impaired downstream Treg responses when the IFN system is blunted. Consistent with these findings, protection against experimental colitis by *B. fragilis* is lost in *Ifnar1*^-/-^ mice. This prompted us to investigate whether tonic IFN induced by commensals poise DCs towards a tolerogenic state. *B. fragilis* treatment of WT DCs led to increased expression of the anti-inflammatory cytokines IL-10 and IL-27. In contrast, *B. fragilis*-treated *Ifnar1*^-/-^ DCs resulted in significantly reduced IL-27, which is known to function downstream of IFNβ signaling (Molle et al., 2007). Of particular interest is the function of IL-27 as an immunoregulatory cytokine that induces IL-10 production to promote development of tolerogenic Tregs (Yoshida and Hunter, 2015; Kim et al., 2019), and suppresses IL-17 producing T cells to prevent inflammation and autoimmunity (Batten et al., 2006; Kim et al., 2019; Nguyen et al., 2019). We also observed a corresponding increase in pro-inflammatory cytokines IL-1β and IL-6 among *Ifnar1*^-/-^ DCs in response to *B. fragilis*. It has been well established that excessive IL-1β and IL-6 leads to the expansion of Th17 cells (Korn et al., 2009). Several studies have demonstrated type I IFN signaling restrains Th17 during autoimmune inflammation and infections (Guo et al., 2008; Prinz et al., 2008; Shinohara et al., 2008). Thus, the reduction in IL-10 and IL-27, along with the pro-inflammatory shift in IFNAR-deficient DCs suggests commensal-derived IFN signaling is essential for tolerogenic DC function and promoting mucosal homeostasis.

Our findings indicate that type I IFN signaling is critical for maintaining immune tolerance. Studies with mice and human CD4+ T cells reported an IFN gene signature in Tregs, corroborating our single cell sequencing data on gut Tregs (Miragaia et al., 2019; Szabo et al., 2019; Sumida et al., 2022). Consistent with this, the locus containing IFNAR, the receptor of type I IFNs, was identified as a susceptibility region for IBD (Jostins et al., 2012). IBD has long been considered the result of loss of immune tolerance for commensal bacteria (Cohen et al., 2019). Further, we reveal that NOD2, a cytosolic pattern recognition receptor in which mutations are strongly associated with IBD (Jostins et al., 2012; Cohen et al., 2019), is required for commensal-derived IFN expression. These studies support the notion that patients with defects in IFN signaling may be susceptible to IBD. Type I IFNs have been investigated for their clinical use in IBD with systemic administration of type I IFNs showing anti-inflammatory effects in both Crohn’s disease and ulcerative colitis. While the effectiveness of type I IFN therapies has been inconsistent in IBD (Nikolaus et al., 2003; Musch et al., 2007), this treatment may selectively benefit those with intact IFN signaling pathways. Moreover, a number of IFNβ drugs are currently approved for the treatment of multiple sclerosis, where outcomes include increased anti-inflammatory cytokine production and reduction in disease relapse (Graber et al., 2007; Cohan et al., 2021).

The link between the microbiota and type I IFN has been well-documented over the last decade. The specific bacterial strains associated with IFN signaling differ amongst these studies. Yet, it is important to note the effect of the microbiota is not dependent on a single species, but rather, a diverse community of microbes can recapitulate type I IFN responses. Our study here supports a model whereby commensal bacteria fine-tune type I IFN to orchestrate immune regulation. While the present study describes a system for the commensal *B. fragilis* to direct immune tolerance in the gut, more broadly our findings can be extended to general mechanisms of immune regulation mediated by the microbiota. For instance, the contribution of the microbiota to responsiveness to immune checkpoint inhibitors (Vétizou et al., 2015; Routy et al., 2018; Matson et al., 2018; Lam et al., 2021) may be linked to the regulation of type I IFN signaling. In line with this notion, *B. fragilis* has been associated with improved response to immune checkpoint blockade in cancer patients (Vétizou et al., 2015) – perhaps owing to its IFN-promoting functions demonstrated here. These observations provide strong evidence that commensal bacteria regulate host immune response through type I IFN pathways.

## Acknowledgements

We thank members of the Chu lab for technical support and helpful discussions. We thank A. Khosravi and G. Sharon for critical reading of the manuscript. We thank the Flow Cytometry Core Facility at the La Jolla Institute and Cheryl Kim for their expertise and instrument support. This work was supported by R00 DK110534 and P30 DK120515 from the National Institute of Diabetes, Digestive and Kidney Diseases (NIDDK) and the Chiba University-UC San Diego Center for Mucosal Immunology, Allergy and Vaccines (cMAV) to H.C. Additional support was provided to H.C. by CIFAR Humans and the Microbiome Program and The Hartwell Foundation.

M.C.T was supported by T32 DK007202 (NIDDK), the National Academies of Sciences, Engineering and Medicine through the Predoctoral Fellowship of the Ford Foundation, and the Howard Hughes Medical Institute (HHMI) Graduate Fellowships grant (GT15123).

## Author Contributions

H.C. conceived the project and designed the experiments. A.V.A., C.Y.H., K.M., E.B., M.C.T., L.R.L., H.H.L. and H.C. performed the experiments. J.H.P. performed the single cell analysis experiments, P.R. and K.M. performed computational analyses, and A.V.A., K.M., C.Y.H., H.H.L. and H.C. analyzed the data. M.T. supported and supervised the single cell RNA sequencing studies. H.C. supervised the project and wrote the manuscript, with contribution from all authors.

## MATERIALS & METHODS

### Mice

Specific pathogen-free wildtype C57BL/6J (Stock No. 000664), *Ifnar1*^-/-^ (Stock No. 028288), *Tlr2*^*-/-*^ (Stock No. 004650), and B6.129-*Ifnβ1*^tm1Lky^ (Stock No. 010818) were purchased from The Jackson Laboratory. *Nod2*^fl/fl^ mice were obtained from G. Nunez (Kim et al., 2016) and crossed with CD11c-Cre mice (Stock No. 008068; The Jackson Laboratory) to generate conditional knockout mice. Mice were maintained on Cre+/-X Cre-/-breeding system. Germ-free C57CL/6J were bred and housed in flexible film isolators. Foxp3^hCD2^/IL-10^Venus^ were obtained from K. Honda (Atarashi et al., 2013) and re-derived germ-free.

For gnotobiotic and mono-colonization experiments, 4- to 6-week-old male and female germ-free mice were orally gavaged with a single dose of *Bacteroides fragilis* (10^8^ CFU resuspended in sterile PBS). Mono-colonized mice were housed in autoclaved cages with autoclaved chow and drinking water supplemented with 100 µg/ml of gentamicin (VetOne). *B. fragilis* strains used are naturally resistant to gentamicin. Fresh fecal samples were collected weekly during cage changing for plating on selective media and germ-free or *B. fragilis* colonization status were confirmed by PCR. Mice were intraperitoneally injected with 100 ug/mouse of polyI:C (Sigma-Aldrich) and housed for 4 hours.

All procedures were performed in accordance with the guidelines and approved protocols from the IACUC of UC San Diego.

### Bacteria

*Bacteroides fragilis* (NCTC9343), *Bacteroides vulgatus* (ATCC 8482), *Anaerostipes caccae* (DSMZ 14662), *Bacteroides thetaiotaomicron* (DSMZ 2079), *Bifidobacterium longum* (NCC 2705), *Blautia producta* (DSMZ 2950), *Clostridium butyricum* (DSMZ 10702), *Clostridium ramosum* (DSMZ 1402), *Escherichia coli* K-12 (MG1655), and *Lactobacillus plantarum* (DSMZ 20174) were grown in Brain Heart Infusion (BHI; Thermo Scientific) media supplemented with 0.5 μg/ml vitamin K (Sigma-Aldrich) and and 0.5 % hemin hemin (Sigma-Aldrich) for 72 hours in anaerobic conditions at 37 ºC. Colonies were inoculated into BHI broth supplemented with vitamin K and hemin to grow overnight in anaerobic conditions (10% H_2_, 10% CO_2_, 80% N2; Coy Lab Products) at 37 ºC. Colonies were inoculated into liquid BHI-S to grow anaerobically for 16-18 hours, then sub-cultured and grown to an OD_600_ of 0.3 (equivalent of 2.1 × 10^8^ CFU/ml) where cells were then heat killed at 90°C for 40 minutes and used for treatment.

### DNBS colitis

WT and *Ifnar1*^-/-^ mice were orally gavaged with PBS or *B. fragilis* (2 × 10^8^ CFU) every other day for one week prior to 2,4-dinitrobenzenesulfonic acid (DNBS; Sigma) administration. On day 7, mice were anesthetized with isofluorane, and rectal administration of 5.5 % DNBS in 50 % ethanol was applied through a 3.5F catheter (Instech Solomon), as previously described (Chu et al., 2016). Control groups received 50 % ethanol (Sham). Briefly, a flexible silicone catheter was inserted 4 cm into the colon, and the mice were held in a vertical position for at least 1 min after rectal administration. Mice were monitored and weighed daily for the duration of the experiment. Upon sacrifice, gut tissue was harvested and colonic lamina propria cells were isolated for further analysis.

### RNA isolation and quantitative real-time RT-PCR

Cells were harvested, washed, and immediately lysed in RLT buffer for RNA isolation using the RNeasy Mini Kit, according to the manufacturer’s protocol (Qiagen). Tissue was harvested and placed in RNA*later* (Invitrogen), homogenized, and lysed according to the manufacturer’s protocol (Qiagen). One ug of RNA was reversed transcribed using SuperScript IV Reverse Transcriptase according to the manufacturer’s protocol (Thermo) and diluted to 10 µg/µl based on the input concentration of total RNA. Gene specific primers were synthesized by Integrated DNA Technologies and sequences are provided in Table 1. Real-time PCR was performed on cDNA using the QuantStudio (Thermo).

**Table 1.**
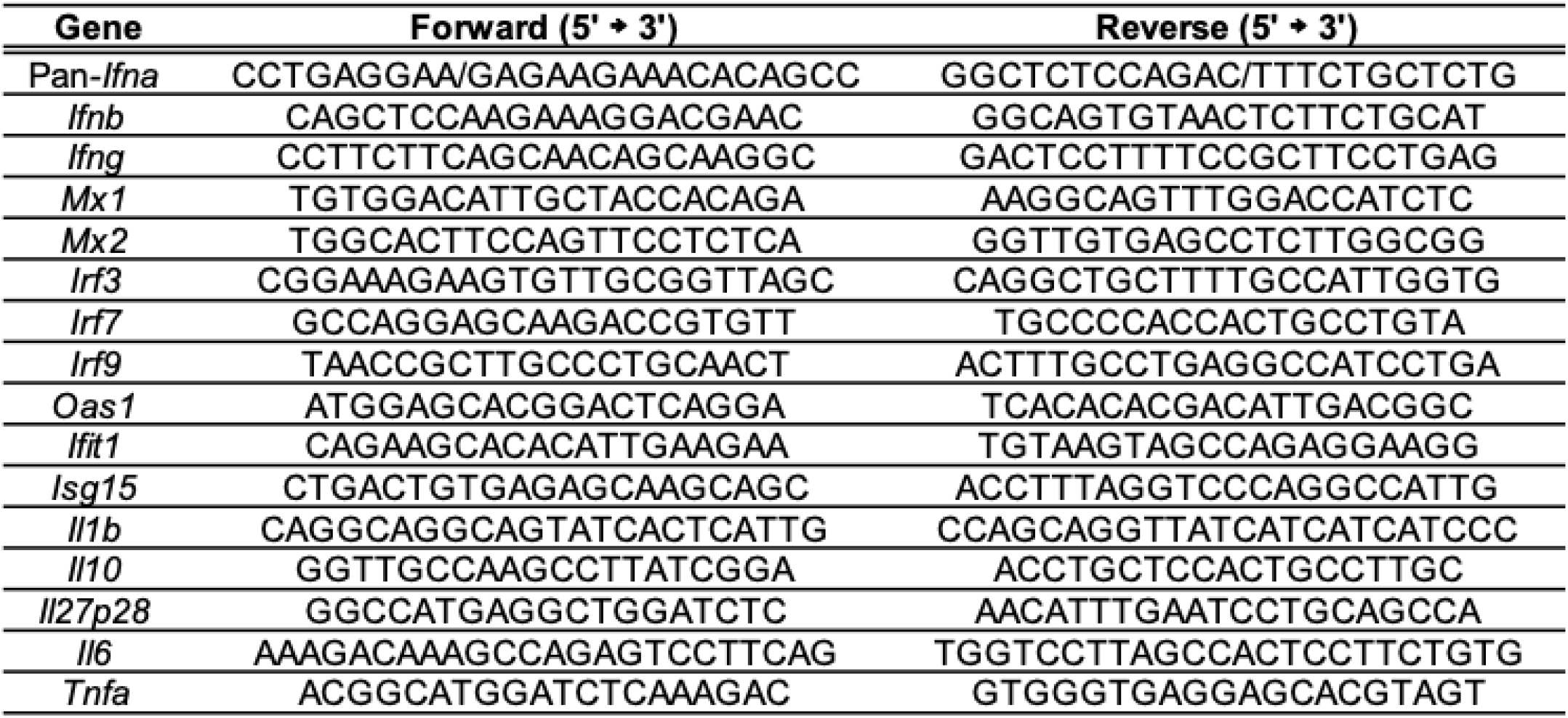
Primers used for real-time PCR.

### Colon explant cultures

1 cm of colon was cut from tissue cleaned with 1X PBS. The tissue was placed in 500 µl of complete RPMI 1640 (10 % fetal bovine serum, 50 U/ml penicillin, 50 µg/ml streptomycin, 2 mM L-glutamine, 1 mM sodium pyruvate, 1 mM HEPES, non-essential amino acids, and beta-mercaptoethanol; Gibco) and cultured for 18-24 hours with or without stimulation with polyinosinic-polycytidylic acid (poly I:C; Sigma) in 37 ºC and 5 % CO_2_. Supernatant were then collected and stored in -20 ºC and used for quantification of cytokine secretion by ELISA.

### *In vitro* DC:T cell co-culture

Co-culture of bone marrow-derived dendritic cells (BMDCs) and CD4+ T cells were performed previously described (Chu et al., 2016). Briefly, BMDCs were generated from bone marrow progenitor cells isolated from femurs of WT or IFNAR1-deficient mice in the presence of 20 ng/ml GM-CSF (Miltenyi) in complete RPMI 1640 (10 % fetal bovine serum, 50 U/ml penicillin, 50 µg/ml streptomycin, 2 mM L-glutamine, 1 mM sodium pyruvate, 1 mM HEPES, non-essential amino acids, and beta-mercaptoethanol). BMDCs were pulsed with PBS, *Bacteroides fragilis* at MOI of 1 or 5 for 18-24 hours. BMDCs were washed and co-cultured with splenic CD4+ T cells at a ratio of 1:10 (DC:CD4+ T cells) in the presence of anti-mouse CD3 (eBiosciences), mouse IL-2 (Peprotech), and human TGF-beta (Peprotech). Co-cultures were treated with 10 µg/ml of anti-Mouse IFNα and anti-Mouse IFNβ (Leinco Technologies) every 24 hours. After 3 days of co-culture, supernatants were collected and stored at -20 ºC for ELISAs and cells stained with specific antibodies and viability dye for analysis by flow cytometry.

### Flow cytometry and ELISA

Cells were stained for 30 min at 4 ºC with either LIVE/DEAD fixable violet or yellow dead stain Kit (Life Technologies), with empirically titrated concentrations of the following antibodies: PE-Cy7-conjugated anti-mouse CD4 (clone: RM4-5), PE-conjugated anti-human CD2 (clone: RPA-2.10), APC-conjugated anti-mouse CD25 (clone: PC61.5), BV785-conjugated CD11c (clone: N418), PE-Cy7-conjugated anti-mouse Siglec H (clone: eBio440c), eF660-conjugated anti-GFP (clone: 5F12.4). For intracellular staining, cells were fixed and permeabilized using the Transcription Factor Phospho Buffer Set (BD Pharmigen) according to the manufacturer’s protocol. Intracellular staining was performed with the following antibodies: AlexaFluor647 - conjugated anti-mouse pSTAT1 (clone: Ser727), PerCP-eF710-conjugated anti-mouse ROR gamma (t) (clone: AFKJS-9), PE-Dazzle594-conjugated anti-mouse Tbet anti-mouse (clone: 4B10), PE-conjugated anti-mouse IL-10 (clone: JES5-16E3), PE-Cy7-conjugated anti-mouse IL-17 (clone: eBio17B7), BV785-conjugated anti-mouse IFNg (clone: XMG1.2), and/or FITC-conjugated anti-mouse Foxp3 (clone: FJK-16s) for 3-4 hours. All antibodies were purchased from Thermo Scientific/eBiosciences, BD, and Biolegend. Cell acquisition was performed on FACSCelesta (BD), and data was analyzed using FlowJo software suite (TreeStar). For ELISAs, cell supernatants from DC or DC-T cell co-cultures were collected and measured using a commercially available kit for IL-10 (Thermo Scientific/eBiosciences), IFNβ (PBL Assay Science), and antiviral response analytes (Biolegend).

### Isolation of cells from tissues

Mesenteric lymph nodes (MLN) were processed by mashing tissues through 100 µm cell strainer (BD Falcon) to generate single cell suspensions. Colon tissues were cut open longitudinally and luminal contents were flushed with cold PBS. Colon tissues were cut into 1 cm pieces and incubated in 10 mM DTT with gentle shaking at 37 ºC for 20 min, followed by incubation in 20 mM EDTA for 20 min. Supernatant were removed and the remaining tissues were incubated in 1 mg/ml Collagenase D and 0.5 mg/ml DNase I. Colonic lamina propria cells were filtered through a 70 µm cell strainer and separated by 40 %:80 % Percoll density gradient. Enrichment of CD4+, CD11c+, and Foxp3-hCD2+ cells were performed using MACS beads (Miltenyi).

### Ex vivo cultures

Lymphocytes isolated from the spleen or colonic lamina propria were cultured in complete RPMI 1640 (10 % fetal bovine serum, 50 U/ml penicillin, 50 µg/ml streptomycin, 2 mM L-glutamine, 1 mM sodium pyruvate, 1 mM HEPES, non-essential amino acids, and beta-mercaptoethanol). Splenocytes were treated with *B. fragilis* at MOI of 10. Colon cells were either untreated or treated with 25 µg/ml recombinant mouse IFN-β1 (Biolegend.) Supernatant was collected for ELISA and cells were stained for flow.

### Single cell RNA sequencing

Mesenteric lymph nodes (MLN) and colonic lamina propria (cLP) single-cell suspensions were generated and approximately 8,000-10,000 cells were loaded onto a Chromium Single Cell Chip (10x Genomics), according to the manufacturer’s protocol. mRNA was barcoded during cDNA synthesis and pooled for Illumina sequencing. Libraries were sequenced with an 8 base index read on a Novaseq 6000. FASTQ files were demultiplexed and aligned using Cell Ranger. Analysis was performed using Loupe (10x Genomics).

### Statistical analysis

Student’s t-test was used for pairwise comparisons. One-way and two-way ANOVA with Post-hoc Tukey test were used for comparisons among one or more groups, respectively, using the GraphPad PRISM software. P-value <0.05 was considered significant.

