## Supplemental Figures for "Commensal bacteria promote type I interferon signaling to maintain immune tolerance"

**Supplemental information**

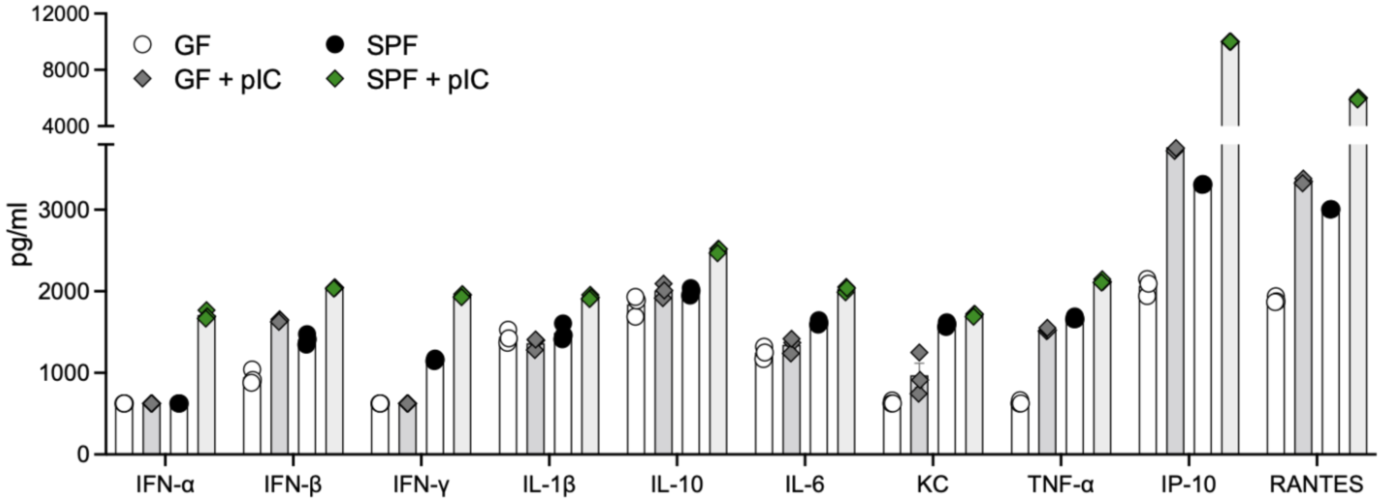


**Figure S1.** GF and SPF splenocytes were treated with 2 µg/ml poly I:C (pIC) for 18 hr. Supernatant was collected to measure cytokine and chemokine secretion by multiplex ELISA. Statistical significance was determined by two-way ANOVA. p<0.05 (*), p<0.001 (***), and p<0.0001 (****).

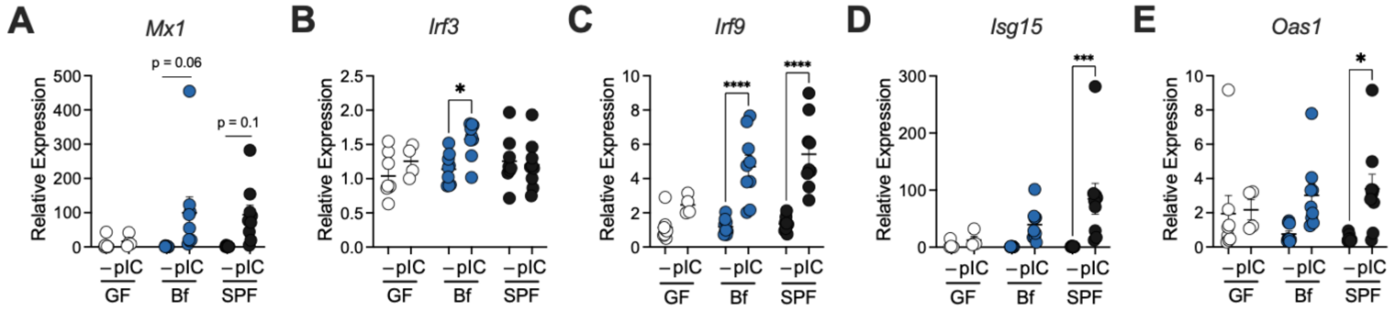


**Figure S2. (A-E)** GF, *B. fragilis* mono-colonized (Bf), and SPF mice were injected (IP) with 100 µg/ml poly I:C (pIC) and colon tissues were harvested after 4 h post-injection. Gene expression for (**A**) *Mx1*, (**B**) *Irf3*, (**C**) *Irf9*, **(D)** *Isg15*, and (**E**) *Oas1* was measured. Each point represents a single mouse. Data are representative of two experiments. Statistical significance was determined by two-way ANOVA. $p<0.05$ (*), $p<0.001$ (***), and $p<0.0001$ (****).

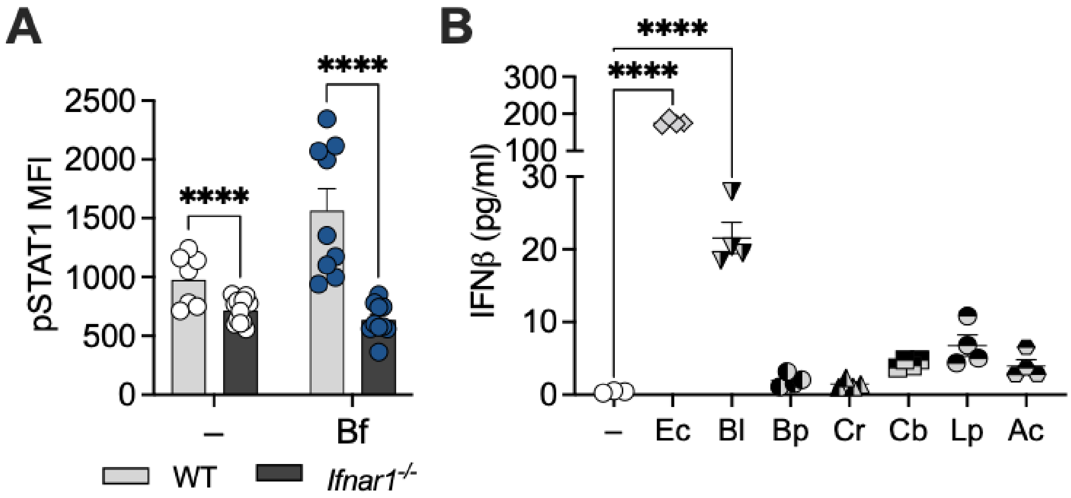


**Figure S3. (A)** WT and IFNAR-deficient BMDCs from SPF mice were treated with *B. fragilis* for 18 hours, and cells were stained for pSTAT1 and analyzed by flow cytometry. (**B**) BMDCs from WT SPF mice were treated with *Escherichia coli*, *Bifidobacterium longum*, *Blautia producta*, *Clostridium butyricum*, *Clostridium ramosum*, *Lactobacillus plantarum*, and *Anaerostipes caccae* for 18 hours. BMDC supernatant was collected and measured for IFNβ by ELISA. Data are representative of two experiments. Statistical analysis was determined by unpaired t-test and two-way ANOVA. $p<0.05$ (*), $p<0.01$ (**), $p<0.001$ (***), and $p<0.0001$ (****).

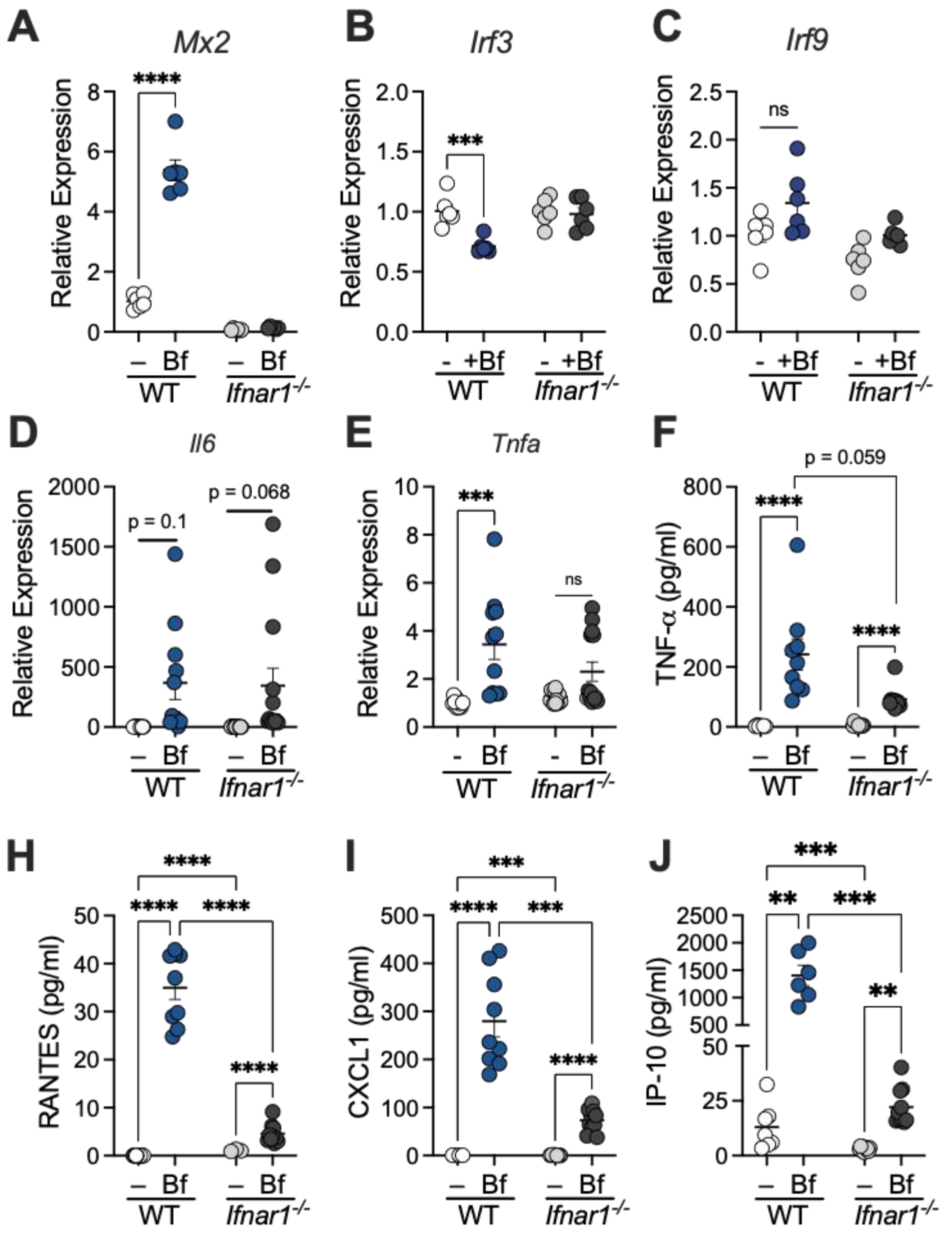


**Figure S4.** WT and IFNAR-deficient BMDCs were pulsed with *B. fragilis* for 18 hours. **(A-E)** Cells were harvested and analyzed by qRT-PCR for expression of **(A)** *Mx2*, **(B)** *Irf3*, **(C)** *Irf9*, **(D)** *Il6*, and **(E)** and *Tnfa* **(F-J)** Supernatant from BMDC cultures were collected and protein secretion was measured for **(F)** TNF-α, **(H)** RANTES, **(I)** CXCL1, and **(J)** IP-10 by LEGENDplex. Data are representative of two experiments. Statistical significance was determined by two-way ANOVA. p<0.01 (**), p<0.001 (***), and p<0.0001 (****).

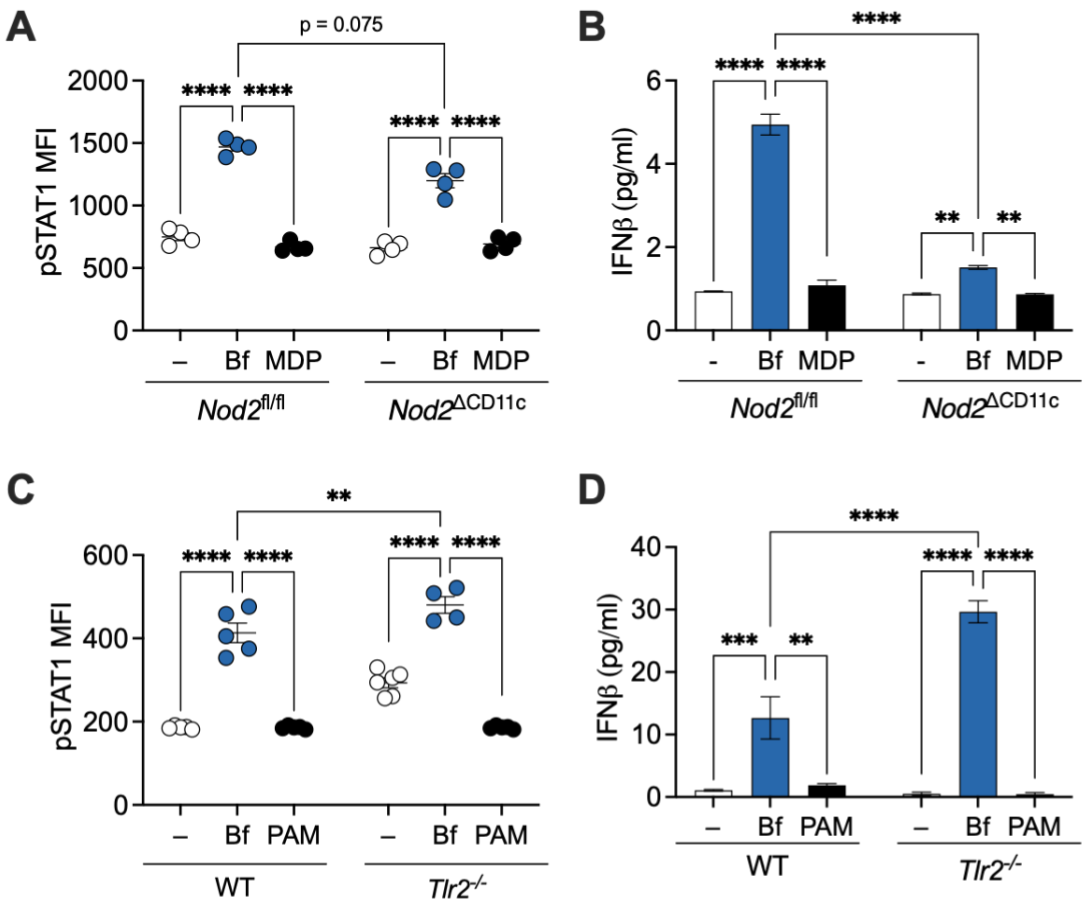


**Figure S5. (A-B)** BMDCs from *Nod2*$^{fl/fl}$ and *Nod2*$^{\Delta CD11c}$ mice were treated with *B. fragilis* or 100 ng/ml of MDP for 18 hours, and (**A**) cells were stained for pSTAT1 and analyzed by flow cytometry. (**B**) Supernatants from BMDC cultures were collected and IFNβ secretion was measured by ELISA. **(C-D)** BMDCs from WT and *Tlr2*$^{-/-}$ mice were treated with *B. fragilis* or 100 ng/ml of Pam3CSK4 (PAM) for 18 hours, and (**C**) cells were stained for pSTAT1 and analyzed by flow cytometry. (**D**) Supernatants from BMDC cultures were collected and IFNβ secretion was measured by ELISA. Data are representative of two experiments. Statistical significance was determined by two-way ANOVA. $p<0.01$ (**), $p<0.001$ (***), and $p<0.0001$ (****).

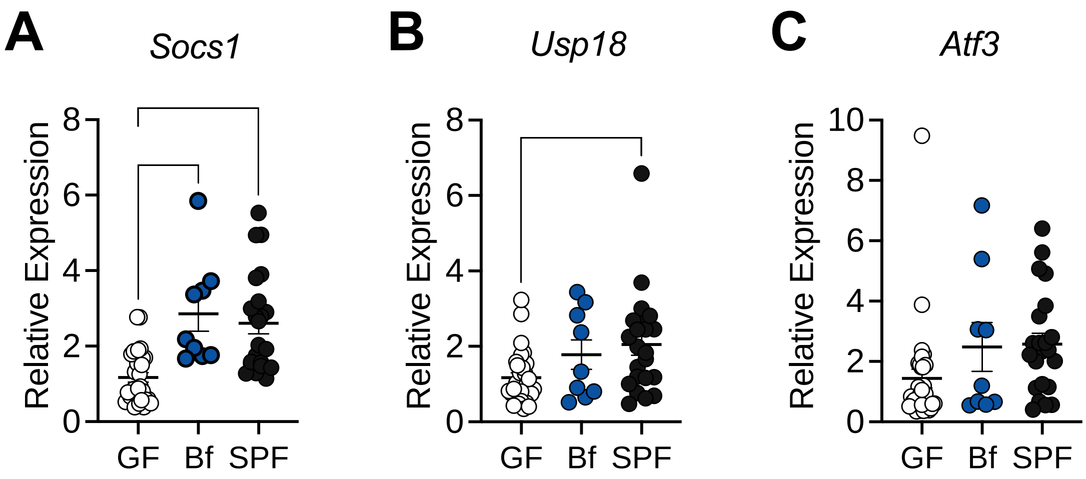


**Figure S6. (A-C)** Colon tissue from germ-free (GF), *B. fragilis* mono-colonized (Bf), and specific pathogen-free (SPF) mice was evaluated by RT-qPCR for expression of **(A)** *Socs1*, **(B)** *Usp18*, and **(C)** *Atf3.* Each point represents a single mouse. Statistical significance was determined by two-way ANOVA. $p<0.01$ (**), $p<0.001$ (***), and $p<0.0001$ (****).

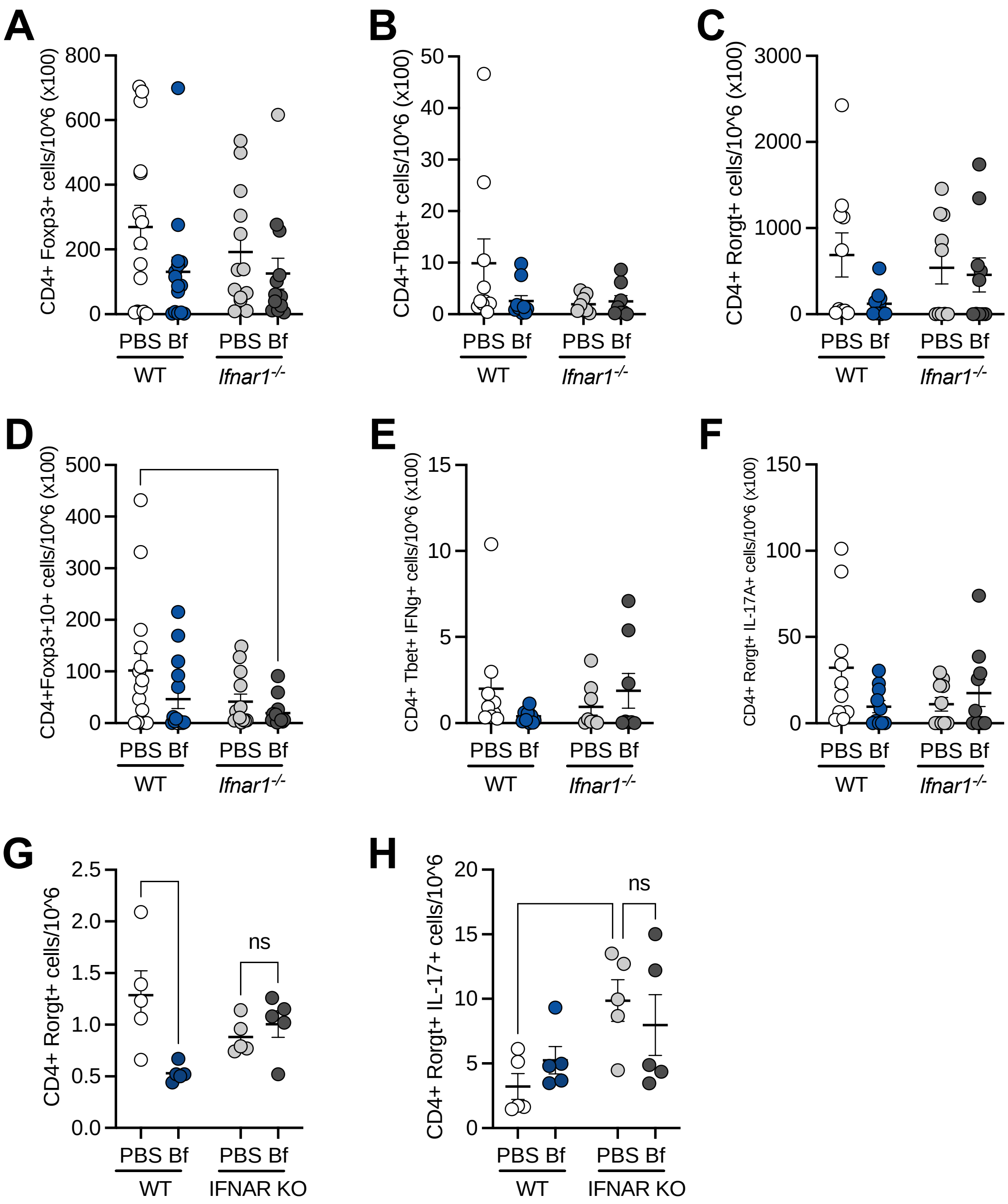


**Figure S7.** WT and IFNAR1-deficient mice were orally treated with sterile PBS or *B. fragilis* during DNBS colitis. Absolute numbers of **(A)** CD4+ Foxp3+, (**B)** CD4+ Foxp3+ IL10A+ (**C)** CD4+ Tbet+, (**D)** CD4+ Tbet+ IFNg+, **(E)** CD4+ Rorgt+, (**F)** CD4+ Rorgt+ IL17A+ T cells in cLP and **(G)** CD4+ Rorgt+, **(H)** CD4+ Rorgt+ IL17A+ T cells in MLN after DNBS colitis were analyzed by flow cytometry. Each point represents a single mouse.
